# Correlated evolution of self and interspecific incompatibility across the range of a Texas wildflower

**DOI:** 10.1101/155986

**Authors:** Federico Roda, Robin Hopkins

## Abstract

Plant species have repeatedly evolved recognition systems between pollen and pistils that identify and reject inappropriate matings. Two of the most important systems recognize self-pollen and interspecific pollen. Outstanding questions are whether and how these two recognition systems are linked and if this association could constrain the evolution of mate choice. Our study characterizes variation in self and interspecific incompatibility in the native range of the Texas wildflower *Phlox drummondii*. We found quantitative variation in self-incompatibility and demonstrate that this variation is significantly correlated with variation in incompatibility with its close congener *P. cuspidata*. Furthermore, we find strong evidence that self and interspecific incompatibility involve common mechanisms of pollen adhesion or early pollen-tube germination. Finally, we show that *P. drummondii* populations that co-occur and hybridize with *P. cuspidata* have significantly higher interspecific incompatibility and self-incompatibility than isolated *P. drummondii* populations. This geographic variation suggests that the evolution of self-compatibility is constrained by selection favoring interspecific incompatibility to reduce maladaptive hybridization. To our knowledge this is the strongest evidence that a correlation between variation in self and interspecific incompatibilities could influence the evolution of pollen recognition across the range of a species.

## Introduction

Mechanisms of mate recognition are among the most rapidly evolving biological systems (Barrett, 2002), from the diversity of floral displays in plants, to the miscellany of mating behaviors in animals. Therefore, a major goal in evolutionary biology is to understand how these mate recognition systems evolve (Stebbins, 1974; Harder & Barrett, 1996; Igic *et al.*, 2008). In plants the most important components of mate choice are whether or not to reproduce with one’s self and whether or not to reproduce with an individual of another species (Barrett, 2002; Barrett *et al.*, 2014). Both of these mate-choices often involve pollen-pistil recognition and rejection systems. There can be significant fitness consequences for allowing or rejecting both self and interspecific pollination. For example, selfing reduces genetic variation and can cause inbreeding depression but might be favored if pollinators or mates are limited and self-pollination results in reproductive assurance (Baker, 1955; Stebbins, 1974; Lloyd, 1992). Conversely, fertilization by interspecific pollen often results in seed abortion, reduced hybrid survival, and hybrid sterility. The potential fitness ramifications for both self and interspecific seed-set establish the importance of understanding if self and interspecific pollen recognition systems share mechanisms and if these common mechanisms constrain the evolution of mate-choice in plants.

The hypothesized link between these two systems is motivated by the correlation between self and interspecific pollen-pistil incompatibility observed in many plants; species that have genetic self-incompatibility (SI) tend to reject pollen of closely related self-compatible species, whereas self-compatible species usually accept interspecific pollen (De Nettancourt, 1977; De Nettancourt, 2001). This asymmetric barrier to hybridization is called “unilateral incompatibility” and has been observed in a variety of genera including *Nicotiana* (Murfett *et al.*, 1996), *Petunia* (Power *et al.*, 1979), *Solanum* (Hardon, 1967), and *Brassica* (Hiscock & Dickinson, 1993b; Schopfer *et al.*, 1999). The pattern suggests the existence of shared mechanisms of self-incompatibility and interspecific incompatibility (II) (Hiscock & Dickinson, 1993a; Hancock *et al.*, 2003a). Molecular support for this link comes from genetic studies in Tomato (De Nettancourt, 2001; Hancock *et al.*, 2003b; Li & Chetelat, 2010; Li & Chetelat, 2014; Li & Chetelat, 2015) and Tobacco (Murfett *et al.*, 1996), where SI genes also control II. However, the ecological consequences of this pleiotropic link remain largely unexplored, in part due to the lack of natural hybridization between species showing unilateral incompatibility and in part due to the lack of genetic variation for the strength of incompatibilities within the species.

Patterns of variation for SI and II within hybridizing species can offer new insights into the forces shaping the evolution of mechanisms of pollen-pistil recognition in plants. For instance, one could explore whether there is a genetic link between self and interspecific pollen recognition mechanisms by testing for a correlation between levels of SI and II across individuals from within a species. Additionally one can determine if the strength of pollen-pistil incompatibilities is greater in areas where the species hybridize than in other parts of the species range. Such pattern could reflect natural selection in the recognition mechanisms to increase reproductive isolation between the species (Levin, 1985; Smith & Rausher, 2007; Smith & Rausher, 2008; Rausher & Bronstein, 2017). Here we addressed these questions using a classic ecological and evolutionary model plant, *Phlox drummondii*. This species segregates genetic variation in both self and interspecific pollen-recognition (Levin, 1985; Bixby & Levin, 1996). Such systems with intraspecific polymorphism in the degree of self-fertilization are particularly valuable to study the forces that influence mating system (Stone *et al.*, 2014; Herman & Schoen, 2016). More importantly, *P. drummondii* co-occur and hybridize with its congener, *Phlox cuspidata*, in a broad area of sympatry in eastern Texas (Levin, 1967; Levin, 1985; Ferguson *et al.*, 1999b). The resulting hybrids are largely sterile (Ruane & Donohue, 2008; Ruane, 2009) indicating selection could favor strong II in sympatry (Levin, 1985; Hopkins *et al.*, 2014).

Our study focuses on variation in SI and II within *P. drummondii*. We first test significant within-species correlation in the strength of SI within *P. drummondii* and II with *P. cuspidata*. We then compare pollen growth and development in self-crosses and interspecific crosses to determine if both types of incompatibilities are generated by similar mechanisms. Finally, we analyze the geographic distribution of incompatibility across the range of *P. drummondii*, to evaluate the impact of environmental variation, in particular *P. cuspidata* presence, in constraining the evolution of these traits. Our study leverages within-species variation in pollen-pistil incompatibility to provide novel insights into if and how self and interspecific mate-choice decisions interact with and constrain evolution of mate-recognition in plants.

## Methods

### Study organisms

*P. drummondii* and *P. cuspidata* are annual wildflowers native to central and eastern Texas. *P. drummondii* has dry stigmas (Heslop-Harrison & Shivanna, 1977) and a gametophytic SI system governed by a major locus (Levin, 1993). However the species shows pseudo-self-fertility (Bixby & Levin, 1996; Levin, 1996b), where the ability of plants to reject their own pollen is often incomplete and varies across individuals and populations (Levin, 1985; Bixby & Levin, 1996; Levin, 1996b; Ruane & Donohue, 2007; Ruane & Donohue, 2008). *P. cuspidata*, is self-compatible and reproduces largely by selfing. These two species diverged around two million years ago (Ferguson *et al.*, 1999a; Ferguson & Jansen, 2002; Roda *et al.*, 2017) and hybridize in areas where they grow together in sympatry (Levin, 1985; Ferguson *et al.*, 1999b; Roda *et al.*, 2017). Both species present similar blue flowers in allopatry but where they come into contact *P. drummondii* has red flower coloration (Levin, 1985). At the edges of these areas of sympatry P. *drummondii* presents strong clines in flower color, with “mixed-color” populations (Hopkins *et al.*, 2014). This flower color variation is the result of reinforcing selection to prevent maladaptive hybridization (Hopkins & Rausher, 2012; Hopkins *et al.*, 2014; Hopkins & Rausher, 2014).

### Population survey

We surveyed variation for SI and II across the natural range of *P. drummondii* (Figure S1, Table S1). For this we collected seeds from natural populations in May of 2014 and 2015, and grew them in the Arnold Arboretum of Harvard University. We sampled 9 *P. drummondii* populations that were sympatric with *P. cuspidata* and 9 allopatric populations. In order to evaluate associations between flower color and pollen-pistil incompatibilities we also sampled 11 populations with mixed flower colors located along color cline transects at the edge of the area of sympatry.

For each population we recorded geographic coordinates with a GPS and collected fruits from ∼ 30 plants located at least 2 m apart (Table S1). To determine if SI variation is explained by mate availability we also measured population density by counting the number of plants on a one square meter quadrat placed at three locations along the most dense parts of the populations (Table S1).

Seeds from the field were stored at 4 C and then grown in HP-mix soil under greenhouse conditions in the absence of pollinators. We obtained an average of 12 ± 3 plants per population, for a total of 343 plants (Table S1).

Because flower color is under selection in *P. drummondii*, we evaluated the possibility of a correlation between reproductive barriers and flower color. For this we qualitatively scored each plant for flower color (light-blue, dark-blue, light-red, or dark-red)(Table S2).

### Seed set analysis

We quantified self-pollination success and interspecific pollination success by measuring seed set after different types of crosses. We emasculated ∼24 flower buds per plant by removing the corollas, which contain the immature anthers. Three days after emasculation, when stigmas were fully developed, we used tweezers to deposit pollen from 2-3 mature anthers onto the stigmas of each emasculated flower. For each plant we conducted three types of crosses: 1) Self-crosses using pollen from the same plant; 2) Interspecific crosses using pollen from a *P. cuspidata* plant; 3) Outcrosses using pollen from a randomly chosen *P. drummondii* plant. Each plant was used once as a pollen-source (Table S2). The outcrosses were used to control for plant maternal fertility and to evaluate baseline levels of reproductive success. It is important to note that we did not find pollination barriers between individuals from different *P. drummondii* populations (data not shown).

We crossed 7 ± 0.15 flowers per plant and cross type (Table S2). Each crossed inflorescence was labeled with tape and bagged with tulle to prevent seed loss after explosive fruit dehiscence. We collected and counted seeds from each crossed inflorescences.

### Pistil observations

To identify developmental mechanisms responsible for pollination barriers we observed *P. drummondii* pistils after self-crosses, out-crosses, and interspecific crosses. In order to capture relevant variation in the strength of pollen pistil-incompatibilities we selected a plants showing broad variation in SI, as determined from seed set. Specifically, we defined plants as plants self-incompatible and self-compatible if they were in the upper and lower 5% tails of the distribution of seed set in self crosses. The three types of crosses were conducted as described previously. We collected pistils 16 hours after pollination and fixed them in FAA (63% ethanol, 5% formaldehyde, 5% Acetic acid). For observation, we washed samples three times with distilled water and then boiled them for 3 minutes in a 5% sodium sulfite solution. Pollen not bound to the stigmas washed away during this process. We then dyed the samples overnight at 4C° in solution of 0.1% Aniline blue in 0.1N Potassium Phosphate buffer. We “squashed” the pistils and observed pollen development and growth using Zeiss Axioskop and Zeiss Axioimager fluorescence microscopes. We observed 6 ± 0.44 pistils per cross-type and maternal sample. For each pistil we counted (1) the number of pollen grains bound to the stigmas, (2) the number of germinated pollen grains, (3) the number of pollen tubes reaching the base of the style and (4) the number of ovules in which the pollen tube penetrated the micropile (Table S3).

Pollen grain counts differ between cross types and phenotypes 16 hours after pollination suggesting that binding is important for pollen-pistil incompatibilities in *P. drummondii* (Figure 2d). We further tested this hypothesis by conducting observations of pollinated stigmas through time, before germination starts (Zinkl *et al.*, 1999). We carried observations of pistils 10 min, 30 min, 1 hour, 2 hours, and 6 hours after pollination for the three types of crosses in self-compatible and self-incompatible plants (Table S3) using pistil squashes as described previously. Additionally we observed the interface between the pollen and the stigma in the Scanning Electron Microscope (SEM) to search for impressions that could evidence a specific molecular interaction between the pollen and the stigma (Zinkl *et al.*, 1999). For this we observed outcross (compatible) pollinations 2 hours after pollination. Fresh samples were cryo-fixed in liquid nitrogen and immediately observed in a JEOL SEM 6010LV SEM.

### Environmental data

In order to identify environmental factors associated with geographic variation in pollen-pistil incompatibilities we quantified environmental variation across. *P. drummondii* populations. We consolidated coordinates from all *Phlox* populations collected in the lab, (Table S4). We then used the geographic coordinates to analyze climatic data from the WorldClim v1.4 database (Hijmans *et al.*, 2005) using the *maptools* (Bivand, 2016), *raster* (Hijmans *et al.*, 2016), *sp* (Pebesma & Bivand, 2005; Bivand *et al.*, 2013) and r*gdal* (Bivand *et al.*, 2016) packages from R. We retrieved monthly measurements of temperature (mean, maximum, minimum) and precipitation as well as altitude and 19 bio-climatic variables using grids with a 30 seconds resolution. We then conducted a principal components analysis (PCA) to separate populations based on all climatic data (Table S4, Figure S2). We used the first three components, which explain most of the variation (PCA1 = 58%, PCA2 = 15%, PCA3 = 6%) to define the climate of the populations. The first PCA largely reflects the latitude (pairwise correlation =-0.87, r-square = 0.76, F[1, 444] = 1437, p < 0.0001), while the second component reflects the longitude (pairwise correlation =-0.89, r-square = 0.80, F[1, 444] = 1833, p < 0.0001).

### Statistical analyses

In this study we wanted to answer three main questions:

1) Is there a correlation between levels of SI and II across *P. drummondii* individuals?
2) What developmental processes explain variation in levels of SI and II?
3) What environmental factors better explain variation in SI and II across *P. drummondii* populations?

To answer these questions we analyzed data from seed counts and pistil observations using generalized linear mixed models (GLMMs). Modeling was conducted with the lme4 package of R (Bates *et al.*, 2014a; Bates *et al.*, 2014b) using a negative-binomial error structure. We included the year of seed collection (i.e. 2014 or 2015) as a fixed effect and the population as a random effect. We tested for over-dispersion in all models and identified significant fixed effects by comparing nested models using likelihood ratio tests. Here we provide a summarized description of the models used. Full models and statistical results are presented in Table S5.

### 1 - Correlations between incompatibilities

We evaluated simultaneously the effect of interspecific seed set, and outcross seed set on self cross seed set.

### 2 - Pollen development

First, we evaluated the effect of cross-type (i.e. self cross, interspecific cross, outcross), plant phenotype (i.e. self-compatible or self-incompatible) and their interaction in the different components of pollen development (i.e. pollen binding, germination, growth and fertilization). Second, we evaluated if there is a correlation between pollination success in self-crosses and interspecific crosses for the different components of pollen development. Third, we evaluated if pollen binding to the stigma increases across time for the three cross types by fitting a linear correlation. Finally we evaluated the effect of the different components of pollen development on seed set.

### 3 - Effect of environmental variables

We explored the effect in seed set from self-crosses and interspecific crosses of the following environmental variables: *P. cuspidata* presence (sympatric / allopatric), geographic location (latitude and longitude), climate (loadings in the first 3 climate PCAs), and population density. Outcross seed set was included in the models as an offset to account for differences in fertility across plants. Because *P. cuspidata* presence is highly correlated with longitude and the second climatic PCA, these three variables were tested in alternative models.

We also evaluated if flower color, which is under divergent natural selection in *P. drummondii*, is correlated with SI and II. Specifically, we evaluated the effect of maternal flower color in seed set from self-crosses and interspecific crosses. For this analysis we used only data from mixed-color populations, where flower color alleles recombine.

## Results

### II and SI are correlated

To estimate the strength of reproductive incompatibilities in *P. drummondii* populations we measured pollen development and seed set in three types of crosses. Self-crosses were used to quantify barriers against self-pollen, interspecific crosses were used to determine interspecific barriers. Reproductive barriers were expected to manifest as reduced seed set when compared to outcrosses between *P. drummondii* individuals.

We found noteworthy variation for seed set in self-crosses and interspecific crosses (Figure 1 a-b). Only 24% of individuals had complete SI (no seeds in self-crosses), and fewer than 10% had complete self-compatibility (equal or greater seed set in self-crosses than outcrosses, Figures 1a and 1c). Interspecific crosses produce significantly more seeds than self-crosses and most *P. drummondii* individuals set fewer seeds when crossed with *P. cuspidata* than with other *P. drummondii* individuals (Figures 1b and 1d).

**Figure 1:**
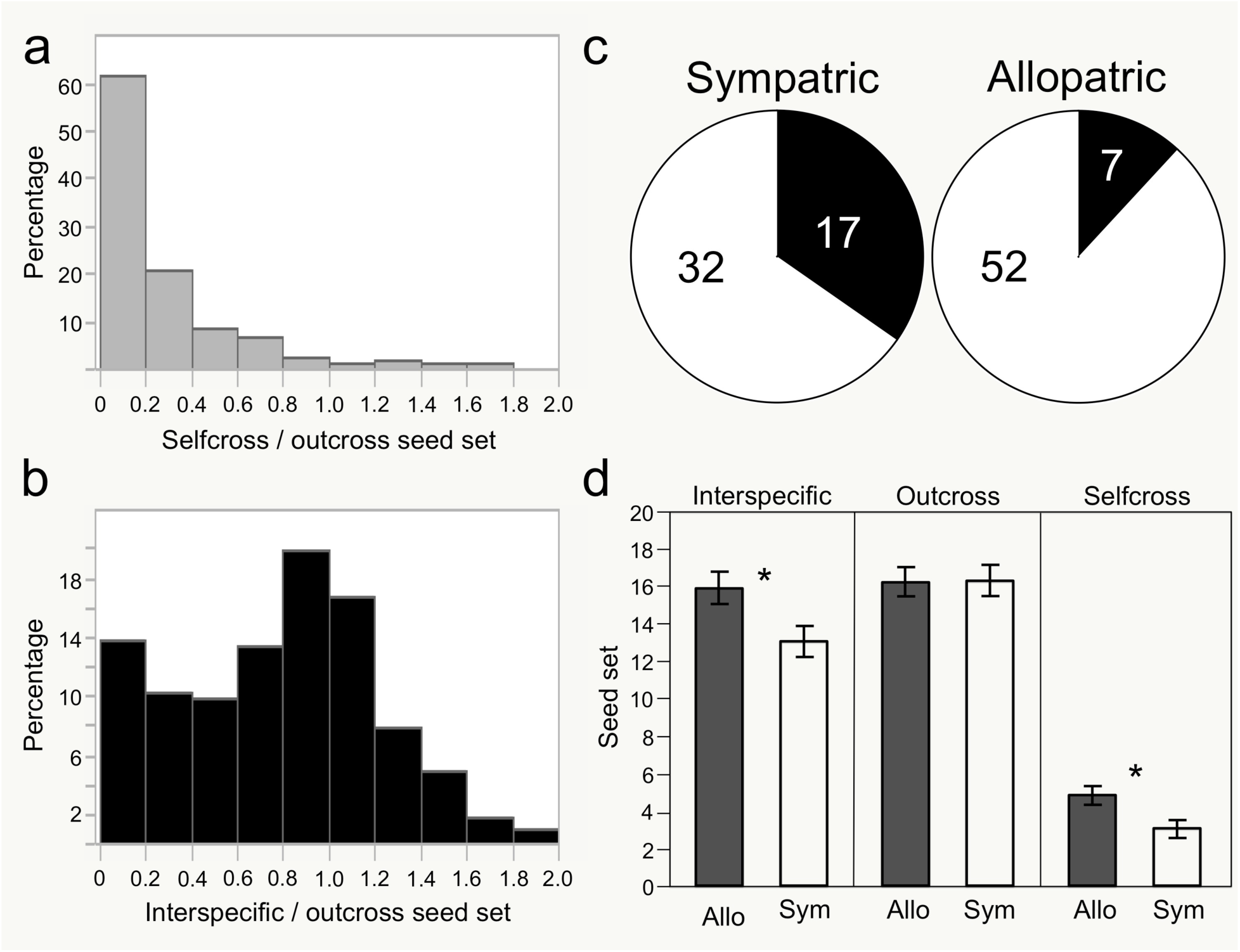
Reproductive barriers are greater in sympatric *P. drummondii* populations. (a-b) Distribution of relative seed set (with respect to outcrosses) in self-crosses (a) and interspecific crosses (b). (c) Number of plants producing seeds in self crosses (white section of pie) in allopatric and sympatric populations. (d) Seed set in allopatric (Allo) or sympatric (Sym) populations after three types of crosses. Means and standard errors are displayed. Asterisks indicate significant differences (p < 0.01) between allopatric and sympatric populations.

**Figure 2:**
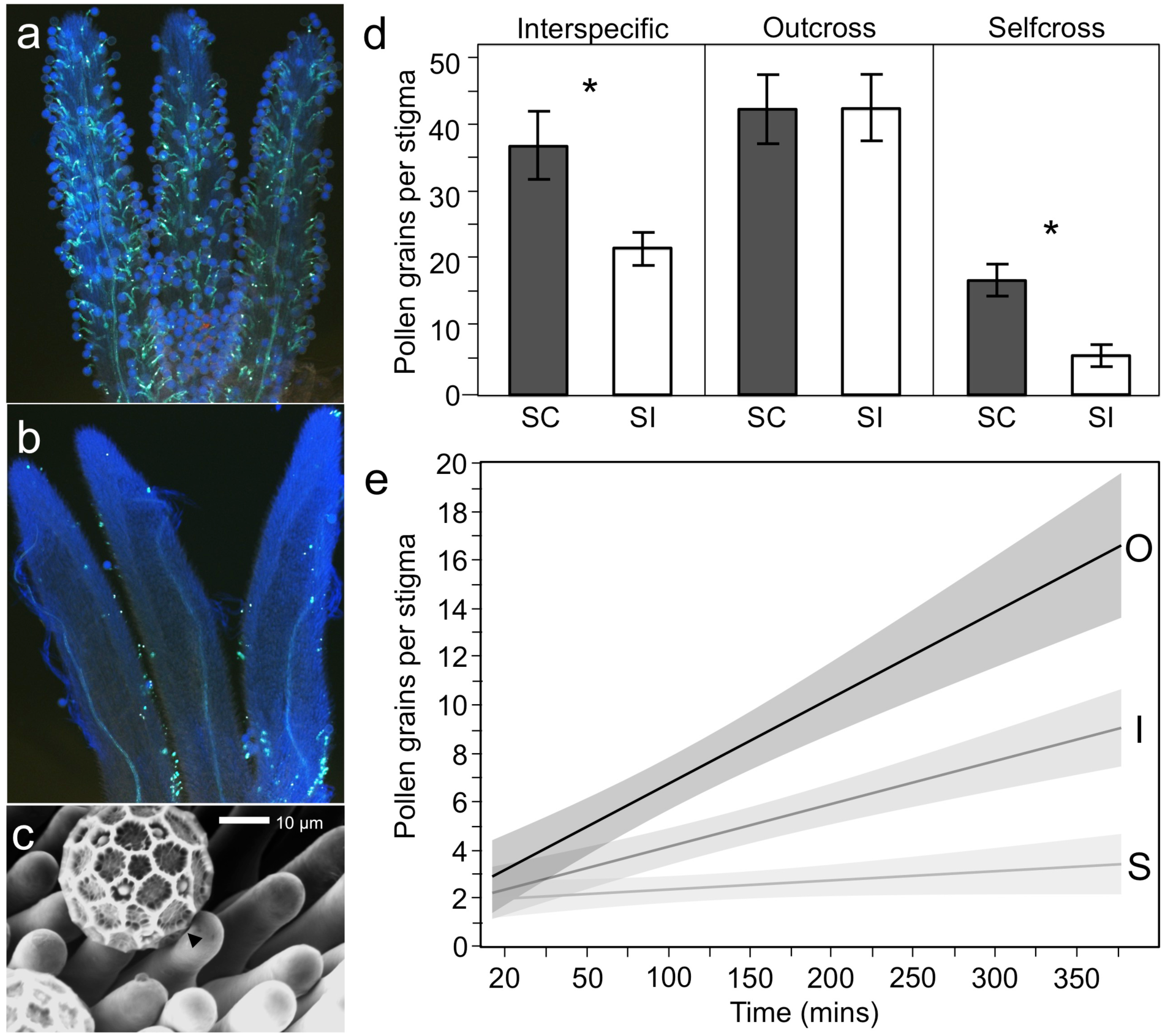
Pollen binding determines reproductive incompatibilities in *P. drummondii:* (a-b) Aniline blue staining of stigmas after compatible (a) and incompatible (b) pollinations. (c) Scanning electron micrographs of pollinated stigmas after outcrosses. Compatible pollen creates impressions in the stigma surface (arrow). (d-e) Assessment of pollen binding to the stigma. Pollen binding in self compatible (SC) and self-incompatible (SI) plants for three types of crosses. Pistils were observed 16 hours after pollination. Means and standard errors are displayed. Asterisks indicate significant differences (p < 0.01) between SI and SC plants. (d) Pollen binding across time. Line of fit showing a linear correlation with confidence intervals for interspecific-crosses (I), outcrosses (O), and self-crosses (S). Means and standard errors are shown in Figure S3.

We used a generalized linear mixed model (Bates *et al.*, 2015) to test if self-pollinated seed set is predicted by interspecific pollinated seed set while controlling for out-crossed seed set and maternal population. Seed set from interspecific pollination significantly predicted self-pollination seed set across individuals, as indicated by a goodness of fit likelihood ratio test (*X*^*2*^(1)= 8.75, p = 0.003. Table S5). This correlation is positive, where individuals that are more self-compatible are also more compatible with *P. cuspidata*.

### Pollen binding is important for SI and II

We further explored the developmental basis of these incompatibilities by observing pollen development in different types of crosses (self-cross, outcross, interspecific cross) and compatibility phenotypes (self-compatible and self-incompatible). We first quantified four different components of pollen development 16 hours after pollination (Figure 2d, Table S5): binding to the papillate stigma, pollen germination, pollen tube growth through the style, and ovule fertilization. We aimed to identify which of these components is involved in incompatible interactions with self-pollen and interspecific pollen.

We found that pollen binding is reduced during self- and interspecific crosses. In general, significantly more outcrossed-pollen adhered to stigmas than self-pollen. Importantly, self-pollen binding was significantly higher in stigmas of self-compatible individuals than in self-incompatible individuals (z-value = 5.46, p < 0.001). Furthermore, similar patterns were observed in interspecific pollinations, such that *P. cuspidata* pollen bound less than outcross-pollen, and *P. cuspidata* pollen binding was significantly higher on self-compatible stigmas than in self-incompatible stigmas (z-value = 3.59, p = 0.004). There is a significant correlation between levels of pollen adhesion for self-crosses and interspecific crosses (*X*^*2*^(1)= 10.38, p = 0.001. Table S5).

In order to confirm that the differences in pollen grain counts 16 hours after pollination are due to differential pollen binding rather than to differences in germination rates we assessed pollen development more rigorously before germination (Zinkl *et al.*, 1999). We first used the SEM to look at the interface between the pollen and the stigma, where binding takes place. Impressions in this interface could evidence alterations in the extracellular protein pellicle or the underlying cell wall (Zinkl *et al.*, 1999). We found that pollen binding leaves impressions in the papillae during compatible interactions (Figure 2c). We also conducted a time-lapse experiment (Figures 2e and S3) to determine if binding increases trough time before the onset of germination, as expected from a specific interaction (Zinkl *et al.*, 1999). We found that pollen binding increases gradually for outcrosses (F[1, 144] = 37.55; p < 0.0001; Rsq = 0.20) and interspecific crosses (F[1, 162] = 49.73; p < 0.0001; Rsq = 0.23) but not for self crosses(F[1, 140] = 2.97; p = 0.0867; Rsq = 0.02). Additionally pre-germination pollen binding was greater in outcrosses than in interspecific crosses and self-crosses (Figure 2e).

Self-crosses have reduced rates of germination, growth and fertilization but these rates did not differ between self-compatible and self-incompatible plants (Table S5). On the other hand interspecific crosses did not show reduced viability for these later components of pollen development (Table S5). When controlling for all other stages of pollination, pollen adhesion is the only trait that significantly predicts seed set across treatments, as evaluated by a goodness of fit likelihood ratio test (*X*^*2*^(1)=4.29, *P*=0.039, Table S5).

### SI and II are greater in sympatric populations

As mentioned previously, there is broad variation in the ability of *P. drummondii* individuals to produce seeds in self-crosses and interspecific crosses. We used GLMMs to evaluate if environmental changes across the range of *P. drummondii* explain this variation in reproductive barriers. We were particularly interested in testing the possibility that SI is affected by the presence of *P. cuspidata* in the eastern part of the range. We also wanted to explore an alternative hypothesis, namely that populations with more limited mate availability have evolved higher self-fertility as a mechanism of reproductive assurance (Busch & Schoen, 2008; Busch & Delph, 2011).

We found that *P. cuspidata* presence has a significant effect on seed set from self-crosses (*X*^*2*^(1)= 5.70, p = 0.017) and interspecific crosses (*X*^*2*^(1) = 6.43, p = 0.011; Table S5). Importantly, sympatric populations have stronger barriers to self and interspecific pollination than allopatric ones (Figure 2, Table S5). Despite this trend there is broad variation in seed set within all populations.

We evaluated climatic variation across populations by conducting a principal components analysis of data reported in the WorldClim database (Figure S2, Table S4). We found that three highly correlated environmental variables showed significant effects on self-compatibility: presence of *P. cuspidata*, geographic longitude (*X*^*2*^(1)= 4.84, p = 0.028) and the second component of climatic PCA (*X*^*2*^(1)= 4.03, p = 0.038). However the model including presence of *P. cuspidata* has a significantly better fit on the data (p < 0.001; Table S5).

We tested two additional hypotheses for the evolution of stronger pollination barriers in sympatric populations. Firstly, we evaluated the possibility of a genetic association between pollen-pistil incompatibilities and flower color, a trait that is under divergent selection in sympatric and allopatric areas (Hopkins & Rausher, 2012; Hopkins & Rausher, 2014). For this we tested for a correlation between flower color and pollination success in mixed color populations, where flower color alleles and self-incompatibility alleles are expected to recombine freely. This analysis showed that flower color has no significant effect on SI (*X*^*2*^(3) = 0.77, p = 0.855) or II (*X*^*2*^(3)= 3.58, p = 0.311). Secondly, we tested for the effect of population density, a proxy for mate availability, in reproductive barriers. Density has no effect on seed set from self-crosses (*X*^*2*^(1)= 0.18, p = 0.696) and interspecific crosses (*X*^*2*^(1)= 0.008, p = 0.977).

## Discussion

Although plants have evolved multiple mechanisms to influence self and interspecific pollination rates, the signaling between reproductive organs is the final and therefore arguably the most important barrier to producing a zygote with a suboptimal mate (Heslop-Harrison, 2000). It has been suggested that SI mechanisms could play an important role in defining plant’s ability to reject interspecific pollen (Hu, 2015; Castillo *et al.*, 2016). Despite being on opposite ends of the genetic-relatedness continuum, Self and interspecific pollen recognition seem to share similar characteristics. Pleiotropic effects of one mate-choice on another could influence the evolutionary trajectory of mate-recognition systems within a species. Here we use a classic system *P. drummondii* wildflowers (Levin, 1985; Levin, 1993; Bixby & Levin, 1996; Hopkins & Rausher, 2012; Hopkins & Rausher, 2014), to test this possibility. We found a correlation between SI and II where both reproductive barriers share developmental pathways and are higher in sympatric populations, suggesting that II imposes a geographic constraint on the evolution of self-compatibility. To our knowledge this is the strongest evidence for a mechanistic link between variation in self and interspecific pollen-recognition within species. We explore how this correlation could influence the evolution of pollen recognition across the range of a species.

### Correlated variation in incompatibilities

The existence of unilateral incompatibility across plant species with different mating systems has motivated the long-standing hypothesis that SI and II have overlapping mechanisms (Lewis, 1958; Abdalla, 1972; Hancock *et al.*, 2003a). However a within-species species correlation between variation in self and interspecific incompatibility has never been demonstrated. Patterns of correlated genetic variation within a species can offer new insights into the evolution of incompatibility and shared molecular mechanisms of self and interspecific incompatibility. In this study we exploited variation in SI to ask if self-incompatible plants are better at rejecting interspecific pollen than self-compatible plants.

The “SIxSC rule” observed in plants species showing unilateral incompatibility predicts that the self-incompatible plant species, *P. drummondii,* will be incompatible with the pollen from the self-compatible species, *P. cuspidata*, while the reciprocal cross won’t show reduced viability. Consistently, our results show that *P. drummondii* sets fewer seeds when pollinated by *P. cuspidata* pollen compared to out-crossed *P. drummondii* pollen (Figure 2). Unpublished results as well as previous reports (Ruane & Donohue, 2007; Ruane & Donohue, 2008) show that *P. cuspidata* does not show barriers against *P. drummondii* pollen.

We found that, although most *P. drummondii* individuals are largely self-incompatible, *P. drummondii* has broad quantitative variation in the strength of SI (Figure 1). Previous studies have shown that this variation is genetically controlled, highly heritable, and amenable to selection (Bixby & Levin, 1996). Here we detected a parallel variation in levels of II across individuals. More importantly, we demonstrate the existence of a within-species correlation of self and interspecific incompatibility, which suggest the existence of common pathways to reject self-pollen and pollen from *P. cuspidata*. The unilateral incompatiblity has been extensively documented in crosses between self-incompatible and self-compatible species (Lewis, 1958; Hancock *et al.*, 2003a; Onus & Pickersgill, 2004; Baek *et al.*, 2015) but, to our knowledge, our results constitute the first report of a correlation of SI and II within individuals of a pseudo-self-fertile species (Baek *et al.*, 2016; Markova *et al.*, 2016; Broz *et al.*, 2017). This correlation between incompatibility mechanisms could result from the sharing of developmental mechanisms of pollen rejection. Alternatively, this pattern could also be explained by the genetic linkage between genetic determinants of both traits or by correlated natural selection on the traits. To begin parsing these options we analyzed pollen development.

### Stigma-pollen binding causes incompatibility

There are many stages at which pollen and pistils can interact to cause incompatibilities (Edlund *et al.*, 2004; Swanson *et al.*, 2004). For a successful pollination pollen must adhere to the stigmatic surface, germinate, grow a pollen tube through the style, find the ovule, and successfully fertilize the egg. Recognition and rejection of self and interspecific pollen can and does occur at any of these stages across flowering plants (Swanson *et al.*, 2004; Iwano & Takayama, 2012; Fujii *et al.*, 2016). Additional research, on new systems is needed to better understand the diversity of developmental mechanisms involved in pollen-pistil recognition systems. We investigated the stage at which incompatibility arises in *P. drummondii* by comparing cross types and incompatibility phenotypes for variation in pollen adhesion, germination, pollen tube growth, and ovule penetration.

In most of plant species SI is controlled by a multi-gene locus called the S-locus that contains both pollen and pistil specificity genes. The evolution of self-compatibility can occur through two mechanisms. Either mutations can “break” the S-locus and allow self-fertilization, or mutations in down-stream interacting molecules can effect the efficiency and function of the SI response (Levin, 1996b; Igic *et al.*, 2008; Raduski *et al.*, 2012). S-locus mutations tend to cause complete self-compatibility (Nasrallah *et al.*, 2002; Okamoto *et al.*, 2007; Boggs *et al.*, 2009; Tsuchimatsu *et al.*, 2010; Raduski *et al.*, 2012). Much less is known about the molecular basis of S-locus independent self-compatibility (but see (Murfett *et al.*, 1996; Tovar-Méndez *et al.*, 2017)). Interestingly, these genetic changes tend to result in pseudo-self-compatibility (Atwood, 1942; Leffel, 1963; Nasrallah, ME & Wallace, DH, 1968; Levin, 1996a). We found reduced rates of pollen binding, germination, growth and fertilization in self-crosses (Figure 2), suggesting that pseudo-self-compatibility in *P. drummondii* is likely mediated by a complex array of pollen-pistil interactions. This is consistent with previous studies indicating that SI in *P. drummondii* is controlled by an S-locus but likely modulated by other genes (Levin, 1993; Bixby & Levin, 1996; Levin, 1996b).

The nature and strength of pollination barriers can differ between types of pollen. For instance, interspecific barriers can involve S-locus dependent mechanisms (Hancock *et al.*, 2003a; Baek *et al.*, 2015; Bedinger *et al.*, 2017) but are often mediated by independent pathways regulating pollen binding (Zinkl *et al.*, 1999; Swanson *et al.*, 2004), pollen tube guidance (Márton *et al.*, 2012) and fertilization (Müller *et al.*, 2016). In the case of *P. drummondii,* we found that the barriers to self-pollen are stronger and more complex than barriers to interspecific pollen (Figures 1d and 2e). However, both types of pollen show reduced adherence to the stigmas (Figure 2), which are dry and papillate (Heslop-Harrison & Shivanna, 1977) and thus require active pollen hydration during adhesion. Our results indicate that the rejection of pollen from both self and interspecific pollination occurs on the stigmatic surface through disruption of pollen adhesion to the stigma. We showed that pollen binding is a progressive and specific process, suggesting that it involves molecular recognition mechanisms between the pollen coat and the papillae (Zinkl *et al.*, 1999; Edlund *et al.*, 2004; Swanson *et al.*, 2004). Regardless of the precise mechanism our analyses indicate that SI and II occur at the same stage of pollen-pistil interaction suggesting similar mechanisms of recognition. Although pollen binding has not been described as a mechanism of gametophytic SI (Clarke & Newbigin, 1993; Golz *et al.*, 1995; Franklin-Tong & Franklin, 2003) it is known to mediate discrimination against self and interspecific pollen in species with dry stigmas and sporophytic SI systems like Arabidopsis (Zinkl *et al.*, 1999; Edlund *et al.*, 2004; Swanson *et al.*, 2004). The possible role of pollen binding in the rejection of unwanted pollen by *P. drummondii* has yet to be explored by quantifying the strength of pollen-pistil adhesion (Zinkl *et al.*, 1999), by manipulating this interaction chemically (Zinkl *et al.*, 1999) or by using molecular genetics to identify the causal genes (Aarts *et al.*, 1993; Aarts *et al.*, 1995; Aarts *et al.*, 1997).

### Geographic pattern of incompatibility suggests evolutionary constraint

Previous research on the interaction between SI and II has focused on the shared genetic basis of incompatibility systems rather than on how and why this correlation evolved (but see (Brandvain & Haig, 2005)). Understanding how shared developmental pathways actually influence the evolution of these traits can provide important insights into the evolutionary causes and consequences of mating system transitions. Transitions between self-compatible and self-incompatible populations often occur in areas of contact between closely related species (Fishman & Wyatt, 1999; Smith & Rausher, 2007; Smith & Rausher, 2008; Grossenbacher & Whittall, 2011; Runquist & Moeller, 2013; Buide *et al.*, 2015), suggesting that interspecific matings can influence the evolution of SI systems (Rathcke, 1983; Campbell & Motten, 1985; Levin, 1985; Stucky, 1985; Randall & Hilu, 1990). Given the strong selection against hybridization between *P. drummondii* and *P. cuspidata* (Hopkins & Rausher, 2012; Hopkins *et al.*, 2014) we hypothesize that there could be selection for II in regions of sympatry to prevent hybrid seed set. This selection could have driven the correlated evolution of higher SI. Consistent with this hypothesis we found that sympatric individuals have significantly lower interspecific seed set than allopatric individuals. Concordantly, we also found that sympatric individuals had significantly lower self-seed set than allopatric individuals. No other environmental or demographic variable has such a strong correlation with SI as *P. cuspidata* presence, suggesting selection on II could be driving the geographic distribution of variation in SI.

This pattern of variation in SI in zones of sympatry is different from other plants, where sympatric populations show higher levels of self-fertility than allopatric ones (Fishman & Wyatt, 1999; Smith & Rausher, 2007; Smith & Rausher, 2008; Grossenbacher & Whittall, 2011; Runquist & Moeller, 2013; Buide *et al.*, 2015). It has been suggested that early autonomous selfing helps these plants to prevent hybridization because self-pollen precedes interspecific pollen in the pistils. Intriguingly, a previous study found that sympatric *Phlox drummondii* populations are more self-fertile than allopatric ones (Levin, 1985), consistently with the pollen precedence hypothesis. We ignore the causes of the discrepancies between these results and the patterns obtained by us, given that the methods used were very similar. We replicated our results across two flowering seasons, which suggest that these where not affected by stochastic variation in self-fertility levels through years.

The correlated expression of multiple characters in sympatric areas can result from independent selection on each character or from selection on one character and correlated responses to selection by the others as a result of pleiotropy or linkage (Levin, 1985). Given that flower color is under divergent natural selection in sympatric and allopatric areas, the observed pattern of variation in mating system could result from linkage between the loci governing flower color and pollen-pistil incompatibilities. However the analysis of mixed color populations showed that seed set does not differ between plants of different colors, suggesting that these two traits are genetically uncoupled and SI evolution is not affected by correlated selection on flower color. On the other hand the correlation between SI and II and the existence of impaired pollen binding in self-crosses and interspecific crosses suggest that II and SI could be controlled by the same genes or by genes participating in the same pathways. Such genetic link between the two pollen recognition mechanisms would constrain the evolution of mating system in the range of *P. drummondii*.

It has been suggested that partial SI expression can represent a “best of both worlds” when pollinators or mates are unreliable, warranting reproductive assurance after opportunities for out-crossing have been exhausted (Nasrallah, M & Wallace, D, 1968; Barrett, 2002; Good-Avila & Stephenson, 2002; Vallejo-Marín & Uyenoyama, 2004; Brennan *et al.*, 2005; Goodwillie *et al.*, 2005; Mable *et al.*, 2005). All *P. drummondii* populations surveyed in this study contain self-fertile plants (Table S2), indicating that polymorphism for mating system is effectively maintained in the populations. Some facts indicate that this variation could provide reproductive assurance while favoring outcrossing. Firstly, outcross pollination success is usually greater than self-pollination success (Figure 1) and self-fertile *P. drummondii* plants can produce seeds in the absence of pollinators by autonomous selfing (Table S2). Secondly, this ability is genetically based and amenable to selection (Levin, 1993; Bixby & Levin, 1996) but can be affected by the environment (Ruane & Donohue, 2007; Ruane & Donohue, 2008) and developmental stage, with older flowers and plants being more self-fertile (Bixby & Levin, 1996). Such systems of delayed selfing are believed to facilitate early outcrossing while providing reproductive assurance (Davis & Delph, 2005; Eckert *et al.*, 2006).

In this context, differences in self-fertility between allopatric and sympatric populations could be reflecting selection for reproductive assurance in allopatric areas due to a lack of pollinators, reduced population sizes, or low plant density (Pannell & Barrett, 1998; Busch & Delph, 2012). However, according to previous studies outbreeding rates in *P. drummondii* populations are independent of population density (Watkins & Levin, 1990). Consistently, we found that population density has no effect on self-seed-set. Although we did not quantify pollinator abundance or population size, field observations suggest that these parameters do not differ between allopatric and sympatric populations. These results indicate that although reproductive assurance could play a role in the maintenance of self-compatible alleles in all populations it likely does not determine variation in self-fertility levels across geography.

Ultimately any test of a hypothesis must include an assessment of alternative hypotheses. For instance, floral manipulations (ie emasculation and pollination exclusion) across the geographical range of *P. drummondii* could allow comparing the magnitude of hybridization, pollen limitation and reproductive assurance between the allopatric and sympatric populations (Kalisz *et al.*, 2004; Smith & Rausher, 2008; Runquist & Moeller, 2013; Barrett *et al.*, 2014; Herman & Schoen, 2016; Layman *et al.*, 2017). Reciprocal transplants between allopatric and sympatric sites could allow dissecting whether differences between regions are caused by adaptation to different pollinator environments (Runquist & Moeller, 2013). Independently of the results obtained in these experiments, we consider that studying the evolution of correlated pollen-pistil recognition systems could enlighten our understanding of the forces influencing the evolution of mating systems in plants.

## Acknowledgements

We thank Juan Losada for his continuous advice on analyses of pollen development. We are thankful to Faye Rosin for her help setting up SEM pictures. We are grateful to Diana Bernal for her contribution in plant collection, rearing and crossing. We are thankful to Stuart Graham for his help during the field trip of 2015. We acknowledge the great help of lab interns Sophie Lattes, Joseph Kearney, Gabriela Garcia, Kattie Cartright, Emily Guo, Sasinat Chindapol, Sarah Gonzalez and Jessica Leslie who helped with plant rearing, seed counting and pistil observations. We thank Callin Switzer for his generous guidance with statistical analyses. We are thankful to all the members of the Hopkins lab for their comments on the manuscript. We thank Kea J Woodruff for greenhouse maintenance.

